# Transport of metformin metabolites by guanidinium exporters of the Small Multidrug Resistance family

**DOI:** 10.1101/2023.08.10.552832

**Authors:** Rachael M. Lucero, Kemal Demirer, Trevor Justin Yeh, Randy B. Stockbridge

## Abstract

Proteins from the Small Multidrug Resistance (SMR) family are frequently associated with horizontally transferred multidrug resistance gene arrays found in bacteria from wastewater and the human-adjacent biosphere. Recent studies suggest that a subset of SMR transporters might participate in metabolism of the common pharmaceutical metformin by bacterial consortia. Here, we show that both genomic and plasmid-associated transporters of the SMR_Gdx_ functional subtype export byproducts of microbial metformin metabolism, with particularly high export efficiency for guanylurea. We use solid supported membrane electrophysiology to evaluate the transport kinetics for guanylurea and native substrate guanidinium by four representative SMR_Gdx_ homologues. Using an internal reference to normalize independent electrophysiology experiments, we show that transport rates are comparable for genomic and plasmid-associated SMR_Gdx_ homologues, and using a proteoliposome-based transport assay, we show that 2 proton:1 substrate transport stoichiometry is maintained. Additional characterization of guanidinium and guanylurea export properties focuses on the structurally characterized homologue, Gdx-Clo, for which we examined the pH dependence and thermodynamics of substrate binding and solved an x-ray crystal structure with guanylurea bound. Together, these experiments contribute in two main ways. By providing the first detailed kinetic examination of the structurally characterized SMR_Gdx_ homologue Gdx-Clo, they provide a functional framework that will inform future mechanistic studies of this model transport protein. Second, this study casts light on a potential role for SMR_Gdx_ transporters in microbial handling of metformin and its microbial metabolic byproducts, providing insight into how native transport physiologies are co-opted to contend with new selective pressures.

**Summary:** Using solid supported membrane electrophysiology, structural biology, and binding assays, we characterize binding and transport of metformin metabolites by bacterial SMR transporters, including proteins associated with horizontal gene transfer in wastewater bacteria that degrade metformin.

## Introduction

Membrane transporters are essential for microbial survival in dynamic environments. They bridge the interior of the cell with the external environment and permit the translocation of nutrients, metabolic byproducts, and toxins across the membrane barrier. In particular, efflux pumps are a first line of defense against a variety of xenobiotics, including anthropogenic chemicals(Kim et al., 2021; Paulsen, 2003). One reflection of the fitness advantage provided by these exporters is their frequent association with horizontal gene transfer (HGT) elements such as integron/integrase sequences and plasmids, which permit useful genes to be shared among bacterial populations. HGT-associated genes encoding drug exporters are especially common among isolates from hospitals, wastewater, agriculture, and other human-adjacent contexts(Pal et al., 2015).

Representatives of the Small Multidrug Resistance (SMR) family of proton-coupled antiporters are among the most common HGT-associated exporters(Pal et al., 2015). These ∼100 residue proteins possess four transmembrane helices per monomer and assemble as antiparallel dimers (Fleishman et al., 2006; Kermani et al., 2022; Kermani et al., 2020). Structures of representative SMRs show a deep aqueous substrate binding pocket with a critical pair of glutamate residues at the bottom(Kermani et al., 2022; Kermani et al., 2020). Substrate and protons compete for binding of these glutamates, ensuring the alternating occupancy inherent to antiport mechanisms(Muth and Schuldiner, 2000). Two SMR subtypes with distinct substrate specificities are commonly associated with HGT (Burata et al., 2022; Kermani et al., 2018; Slipski et al., 2020). These are termed SMR_Gdx_ (**g**uani**d**inium e**x**port) and SMR_Qac_ (**q**uaternary **a**mmonium **c**ation). The SMR_Qac_ proteins are promiscuous exporters of polyaromatic and quaternary ammonium antimicrobials, including common household and hospital antiseptics such as benzalkonium(Saleh et al., 2018; Yerushalmi et al., 1995). Quaternary ammonium antiseptics are one of the original modern antimicrobials, commonly used since the 1930s. The SMR_Qacs_ are perhaps the first, and remain among the most common, HGT-associated efflux pumps(Gillings et al., 2008; Zhu et al., 2017). In contrast, the rationale for widespread association between HGT elements and the SMR_Gdx_ is not as obvious. In their major physiological context, SMR_Gdx_ export the nitrogenous waste product guanidinium (Gdm^+^)(Kermani et al., 2018; Nelson et al., 2017), a compound that is widespread in microbial metabolism(Breaker et al., 2017; Funck et al., 2022; Schneider et al., 2020; Sinn et al., 2021; Wang et al., 2021). The SMR_Gdx_ do not provide robust resistance to classical antimicrobials or antiseptics(Chung and Saier, 2002; Kermani et al., 2018). However, an emerging body of literature suggests that even pharmaceuticals that are not used explicitly as antimicrobials also shape bacterial communities in the human microbiome and other human-associated environments(Maier et al., 2018).

One such pharmaceutical is the biguanide antidiabetic metformin. The most frequently prescribed drug worldwide, over 150 million patients are prescribed metformin annually to manage type II diabetes(Lunger et al., 2017). Metformin is typically dosed in gram quantities daily, and is excreted in an unaltered form(Corcoran and Jacobs, 2022; Gong et al., 2012). Metformin and its associated degradation product guanylurea are the most prevalent anthropogenic chemicals in wastewater globally. Concentrations have been measured up to the low μM range in sampled waste and surface waters, and these compounds are not removed through typical wastewater treatment protocols(Briones et al., 2016; Golovko et al., 2021). As a result, these compounds have accumulated to levels of environmental concern in surface water worldwide(Balakrishnan et al., 2022; Briones et al., 2016; Elizalde-Velazquez and Gomez-Olivan, 2020; Scheurer et al., 2012). Metformin is also associated with changes in the composition of microbial communities including the gut microbiome (Vich Vila et al., 2020; Wu et al., 2017) and in wastewater treatment plants(Briones et al., 2016). In some cases, metformin may act as a co-selective agent, enhancing the survival of antibiotic-resistant bacteria in the presence of antibiotics(Wei et al., 2022). However, other recent studies have isolated bacteria that utilize metformin as a nitrogen and/or carbon source(Chaignaud et al., 2022; Li et al., 2023; Martinez-Vaz et al., 2022), suggesting that biodegradation of metformin and guanylurea may be a viable strategy for remediation of these compounds.

Studies on metformin degradation by microbial communities suggests that SMR transporters might have an emerging role in metformin biodegradation. For example, two identical, adjacent open reading frames encoding an SMR_Gdx_ protein were identified on the same plasmid as other genes that contribute to metformin degradation by a wastewater treatment plant isolate(Martinez-Vaz et al., 2022). We previously showed that this protein possesses guanylurea transport activity(Martinez-Vaz et al., 2022). In an independent study, a transcriptional analysis of a metformin-degrading *Aminobacter* strain showed a 30-fold increase in gene expression of an SMR_Gdx_ transporter in metformin-grown cells(Li et al., 2023). On the basis of these studies, pathways for the full breakdown of metformin by bacterial consortia have been proposed. In such pathways, SMR_Gdx_ transporters would provide a key step in the process, export of the intermediate guanylurea (**Figure 1A**).

**Figure 1.**
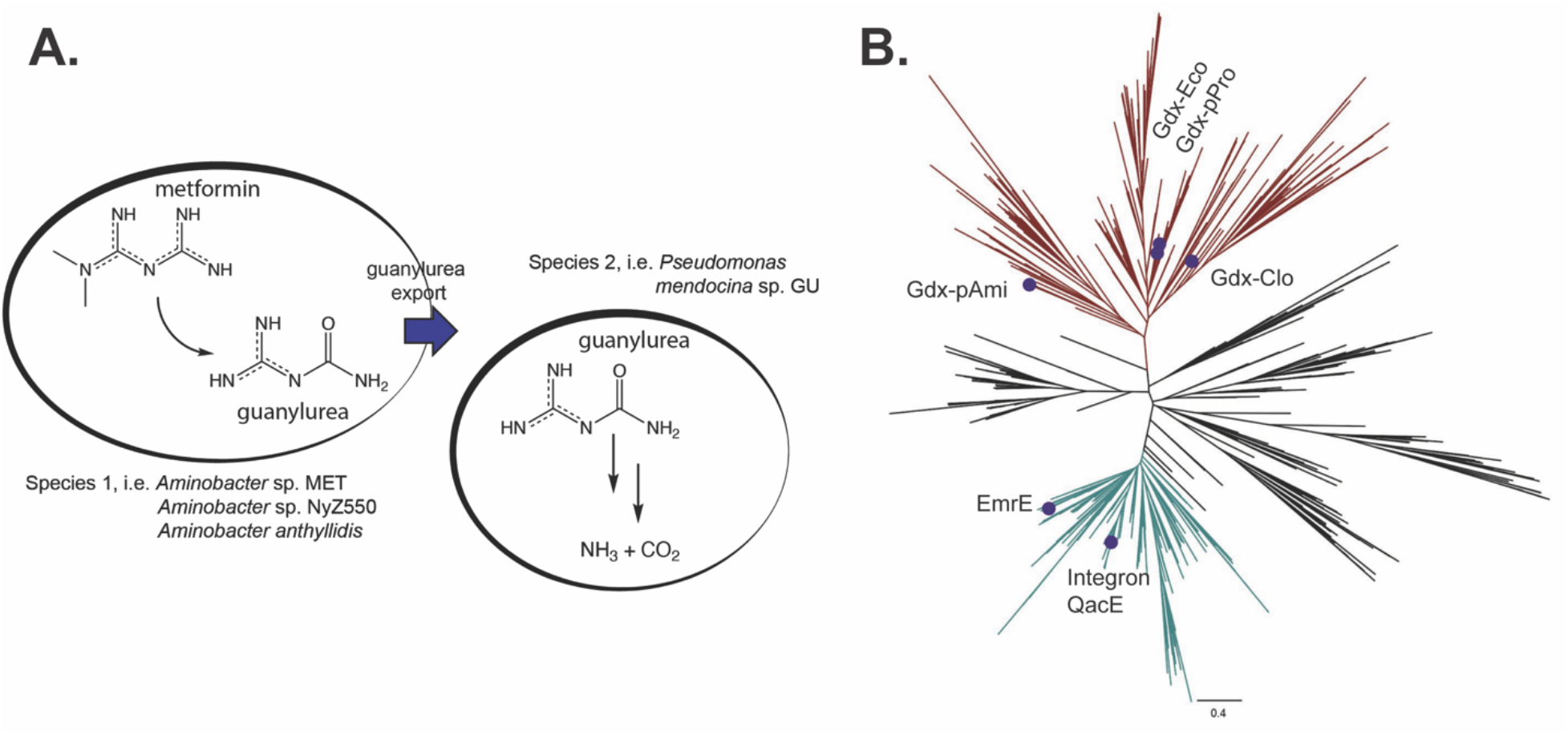
SMR physiology and phylogenetic distribution. A. Schematic showing hypothesized role for horizontally transferred SMR_Gdx_ homologues in biodegradation of metformin by bacterial consortia. B. Phylogeny of the SMR family. SMR_Gdx_ are shown in rust and SMR_Qac_ in teal. Proteins examined in this study are indicated.

In this paper we investigate whether several genomic and plasmid-associated SMRs (**Figure 1B, Supplementary Table 1**) transport metformin or other byproducts of microbial metformin metabolism. For our initial screen, we examined four SMR_Gdx_ homologues and two SMR_Qac_ homologues. The SMR_Gdx_ homologues we examined include 1) the structurally characterized genomic protein from *Clostridales* oral taxon 876, Gdx-Clo(Kermani et al., 2020); 2) the genomic *Escherichia coli* homologue Gdx-Eco(Kermani et al., 2018); 3) a common plasmid-borne variant isolated from multiple species of γ-Proteobacteria, Gdx-pPro(Slipski et al., 2020), which shares 81% sequence identity with Gdx-Eco; and 4) a plasmid-borne variant isolated from *Aminobacter* sp. MET, which uses metformin as a sole nitrogen source, Gdx-pAmi(Martinez-Vaz et al., 2022). We also selected two representatives of the SMR_Qac_ subtype; exemplar EmrE from *E. coli;* and QacE, the most common integron– and plasmid-associated sequence(Burata et al., 2022). We show that efficient guanylurea transport is a general property of the SMR_Gdx_ subtype, but not of SMR_Qac_, and that other metformin degradation products are also transported by SMR_Gdx_. We characterize the transport kinetics and proton-coupling stoichiometry of a representative plasmid-borne and genomic SMR_Gdx_, and determine a structure of a representative SMR_Gdx_ with guanylurea bound. This work provides a case study into bacterial co-option of existing metabolic transporters to deal with novel xenobiotics. Furthermore, this study provides foundational biochemical characterization of the SMR_Gdx_ subtype, which will support future efforts to understand detailed molecular mechanisms of substrate transport by this family of proteins.

## Materials and Methods

### Phylogeny preparation

SMR sequences from representative genomes and from Integrall(Moura et al., 2009), a database of integron-associated genes, were aligned using MUSCLE(Edgar, 2004). A phylogeny was constructed using PhyMl3.0(Guindon et al., 2010) and visualized using FigTree (http://tree.bio.ed.ac.uk/software/figtree).

### Transporter Expression, Purification, and Reconstitution

Gdx-Clo(Kermani et al., 2018), Gdx-Eco(Kermani et al., 2018), EmrE(Kermani et al., 2022), and Gdx-pAmi(Martinez-Vaz et al., 2022) construct design and purification have been described previously. For QacE and Gdx-pPro, synthetic geneblocks (Integrated DNA Technologies, Coralville, IA) were cloned into a pET21b vector with an N-terminal hexahistidine tag and LysC and thrombin recognition sequences. Proteins were overexpressed in C41(DE3). Expression was induced by addition of 0.2 mM IPTG for 3 hours. Cells were lysed and extracted with 2% n-decyl-β-D-maltoside (DM) for 2 hours. After pelleting insoluble cell debris, proteins were purified using cobalt affinity resin. Wash buffer contained 25 mM Tris, pH 8.5, 150 mM NaCl, 5 mM DM. For Gdx-pAmi, NaCl concentration was increased to 500 mM NaCl. The affinity column was washed with wash buffer, then wash buffer with 10 mM imidazole, prior to elution with wash buffer with 400 mM imidazole. For Gdx-Clo and Gdx-Eco, histidine tags were cleaved with LysC (200 ng/mg of protein; two hours at room temperature; New England Biolabs), and for all others, histidine tags were cleaved with thrombin (1 U per mg of protein, overnight at room temperature; MilliporeSigma, Burlington, MA, United States). Proteins were further purified using a gel filtration Superdex200 column (Cytiva, Marlborough, MA) equilibrated with 100 mM NaCl, 10 mM N-2-hydroxyethylpiperazine-N’-2-ethanesulfonic acid (HEPES) pH 7.5, 5 mM DM. Purified proteins were stored at 4° C for up to five days before detergent binding assays. To prepare proteoliposomes for electrophysiology assays, purified protein was mixed with *E. coli* polar lipid extract (10 mg/mL; Avanti Polar Lipids, Alabaster, AL) solubilized with 35 mM 3-[(3-cholamidopropyl)dimethylammonio]-1-propanesulfonate (CHAPS) at a protein to lipid ratio of 40 μg SMR transporter: mg lipid (1:370 protein:lipid molar ratio) prior to detergent removal by dialysis. For preparations that included Fluc-Bpe, liposomes were reconstituted with a molar ratio of 0.3 Fluc-Bpe:1 SMR_Gdx_: 5920 lipid (1 μg Fluc-Bpe and ∼2.5 μg SMR_Gdx_ per mg lipid). For liposome transport assays, proteoliposomes were prepared similarly, except that a 2:1 mixture of 1-palmitoyl, 2-oleoylphosphatidylethanolamine (POPE) and 1-palmitoyl, 2-oleoylphosphatidylglycerol (POPG) (10 mg/mL; Avanti Polar Lipids) was used with 0.2μg protein/mg lipid. Proteoliposomes were stored at –80° C until use.

### Solid supported membrane electrophysiology

SSM experiments were performed using SURFE^2^R N1 instrument (Nanion Technologies, Munich, Germany). Sensors were prepared with a 1,2-diphytanoyl-sn-glycero-3-phosphocholine (DPhPC) lipid monolayer according to published protocols(Bazzone et al., 2017). Each sensor’s capacitance and conductance were verified before use (<80 nF capacitance, <50 nS conductance) using Nanion software protocols. Proteoliposome stock was diluted 1:25 in assay buffer (100 mM KCl, 100 mM KPO_4_ pH 7.5) prior to fusion with the DPhPC monolayer. For substrate screening experiments, positive reference samples were checked periodically; if the current amplitude of the reference compound differed by more than 10% on one sensor, this indicated bilayer instability, and the sensor was not used for further experiments.

*Pyranine Stoichiometry Assay:* Proteoliposomes were reconstituted with an internal buffer of 25 mM HEPES, pH 7.53, 100 mM NaCl, 100 mM KCl and pre-loaded with 0.4 mM substrate (Gdm^+^ or guanylurea) and 1mM pyranine (trisodium 8-hydroxypyrene-1, 3, 6-trisulfonate; Sigma-Aldrich) using three freeze/thaw cycles. Unilamellar liposomes were formed by extrusion through a 400 nm membrane filter and the external pyranine was removed by passing liposomes through a Sephadex G-50 column spin column equilibrated in internal buffer with substrate. The external assay buffers contained 25 mM HEPES, pH 7.53, 0.4 mM substrate, and varying KCl concentration (3-46 mM) to establish the membrane potential, with NaCl to bring the total salt concentration to 200 mM. Proteoliposomes were diluted 200-fold into the external buffer, and after ∼30 s. to establish a baseline, valinomycin (final concentration 0.2 ng/mL) was added together with substrate (final concentration 4 mM). Fluorescence spectra were monitored (._ex_ = 455 nm;._em_= 515 nm) for ∼300 s. The membrane potential was calculated using the Nernst potential for K^+^:

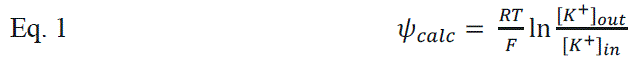

Fluorescence emission timecourses were corrected for baseline drift measured prior to substrate and valinomycin addition. The stoichiometry was determined from the voltage at which electrochemical equilibrium occurred (no change in fluorescence) using the following equation:

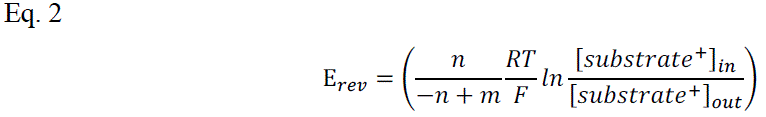

where *n* and *m* represent the stoichiometric coefficients of protons and substrate, respectively.

*Tryptophan Fluorescence:* Fluorescence emission spectra (._ex_ = 280 nm,. _ex_ = 300-400 nm) were collected for 1µM purified protein in assay buffer containing 200 mM NaCl, 10 mM HEPES, 10 mM bicine, 10 mM NaPO_4_, 5mM DM, with pH adjusted from 6.5-9.0. Substrate was added from a stock solution prepared in assay buffer. For Gdm^+^ titrations, the change in fluorescence, *F*, as a function of substrate fit to a single site binding isotherm,

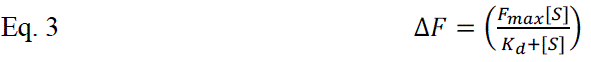

For guanylurea titration, binding data fit to a single site binding isotherm with a correction for a linear, non-specific binding component, *c*:

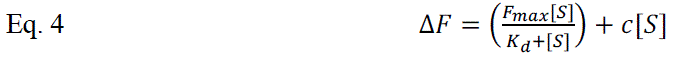

To derive the K_a_ values and K_d_ values from the apparent K_d_ measured as a function of pH, we used the following equation, which uses the approximation that the protonatable E13 sidechains have equal K_a_ values:

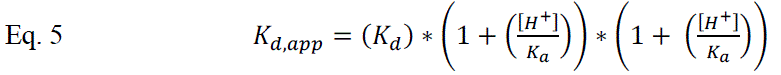

*Isothermal Titration Calorimetry (ITC):* ITC experiments were conducted using a low volume Nano ITC instrument (TA Instruments, New Castle, DE). Freshly purified protein (650 µM) in 10 mM 4-(2-Hydroxyethyl)-1-piperazinepropanesulfonic acid (EPPS), pH 8.53, 100 mM NaCl, 4 mM DM was titrated with 20 mM Gdm^+^ or 10 mM guanylurea prepared in the same buffer. Data was analyzed using NanoAnalyze software.

*Structure of Gdx-Clo in complex with guanylurea:* The crystallization chaperone monobody L10 was prepared as described previously(Kermani et al., 2022; Kermani et al., 2020). Freshly purified Gdx-Clo (10mg/mL) and L10 monobody (10mg/mL, supplemented with 4 mM DM) were mixed at a 1:1 ratio. Guanylurea and lauryldimethylamine-N-Oxide (LDAO, Anatrace) were added to a final concentration of 10 mM and 6.6 mM, respectively, and combined in a 1:1 ratio with crystallization solution. Crystals formed at room temperature after ∼ 7 days in 0.1 M HEPES pH 7.0, 0.1 M calcium acetate, 31% PEG600. Data was collected at the Life Sciences Collaborative Access Team at the Advanced Photon Source, Argonne National Laboratory. Data was processed using DIALS(Winter et al., 2018) software and subjected to anisotropic truncation using Staraniso(Tickle, 2018). Phaser(McCoy et al., 2007) was used for molecular replacement with Gdx-Clo and L10 monobodies (PDB:6WK9) as search models. Coot(Emsley et al., 2010) and Phenix(Liebschner et al., 2019) were used for iterative rounds of model building and refinement.

## Results

### Guanylurea transport is general among SMR_Gdx_ homologues

We first sought to determine whether transport of guanylurea is widespread among SMR homologues, and whether other metformin metabolites might also be exported by transporters from this family. We selected several SMRs that could be purified with monodispersed size exclusion chromatograms (**Supplementary** Figure 1), including both genomic and plasmid-associated SMR_Qac_ and SMR_Gdx_ representatives (see **Figure 1)**. We screened a series of metformin metabolites for transport using solid-supported membrane (SSM) electrophysiology (**Figure 2**). For these experiments, purified proteins are reconstituted into proteoliposomes, which are then capacitively coupled to an electrode to monitor charge movement across the liposome membrane(Bazzone et al., 2017). Because of their antiparallel topology, homodimeric SMR transporters possess two-fold symmetry with identical inward– and outward-facing structures(Morrison et al., 2012); thus, in contrast to most transporters, orienting the proteins in the reconstituted liposome system is not necessary. All compounds were tested for transport at 2 mM, and for each substrate, we confirmed that protein-free liposomes did not exhibit pronounced currents (**Supplementary** Figure 2). Since the efficiency of proteoliposome fusion to the sensors is variable, we included a positive control compound to benchmark the currents for test substrates evaluated on the same sensor: Gdm^+^ for SMR_Gdx_ proteins, and tetrapropylammonium (TPA^+^) for SMR_Qac_ proteins.

**Figure 2.**
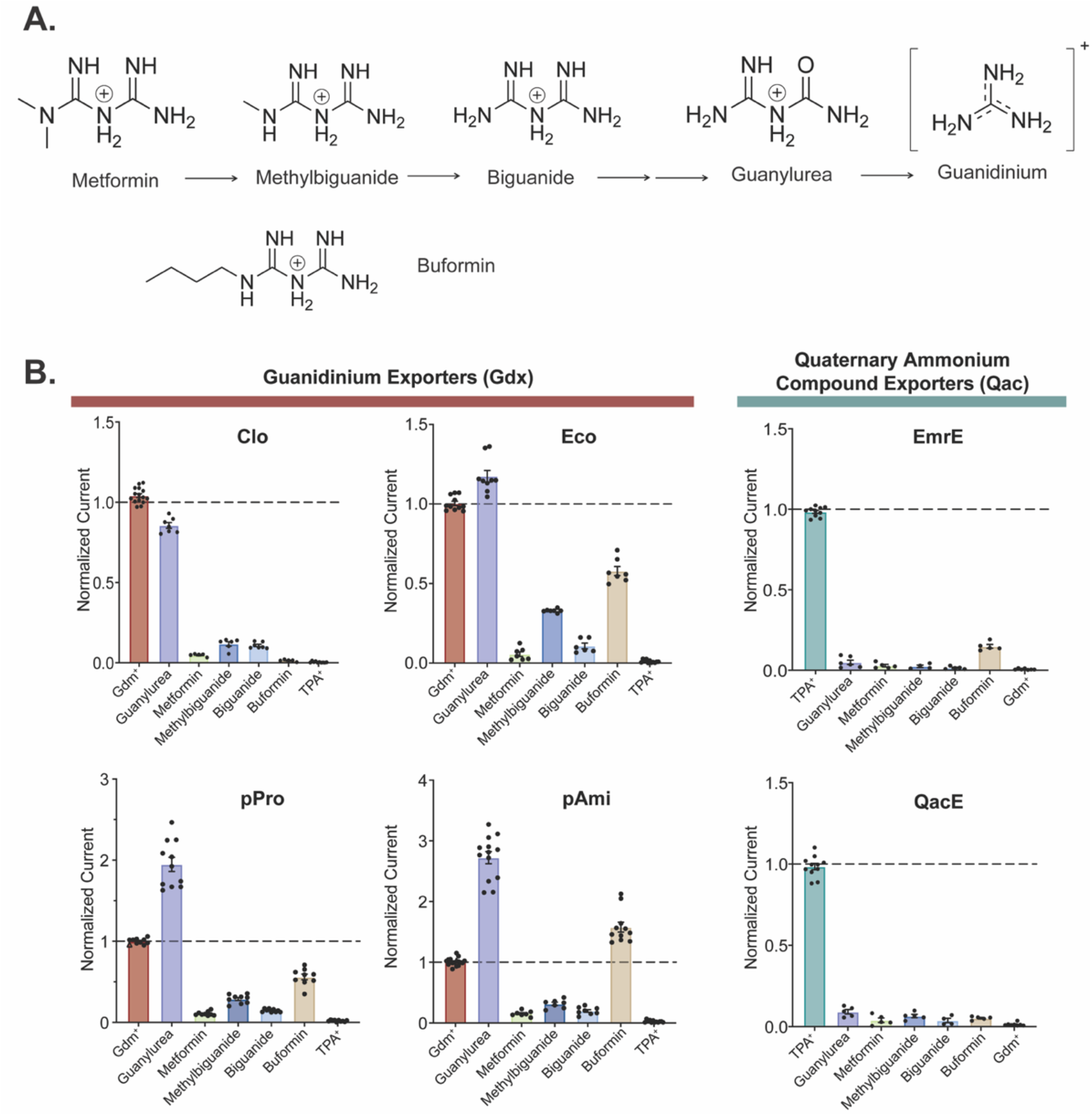
Screen for transport of metformin metabolites by SMR homologues. A. Chemical structures of metformin metabolites and metformin analog buformin. B. Amplitude of transport currents evoked by perfusion with 2 mM substrate. Current amplitudes are normalized to a positive control (Gdm^+^ for SMR_Gdx_ and TPA^+^ for SMR_Qac_) collected on the same sensor. Datapoints represent at least three independent sensor preparations from at least two independent biochemical purifications.

For all SMR_Gdx_ homologs, we observed negative capacitive currents for both Gdm^+^ and guanylurea, consistent with electrogenic proton-coupled substrate antiport (**Supplementary** Figure 2). The best characterized SMR_Gdx_ homologue, Gdx-Clo, transported only Gdm^+^ and guanylurea. However, the other three SMR_Gdx_ homologues tested also transported singly-substituted biguanides, including the metformin degradation product methylbiguanide and the related antidiabetic drug buformin. Metformin, a doubly-substituted biguanide, exhibited currents barely above the detectable limit by SMR_Gdx_ proteins. These observations are congruent with prior observations that guanidinium ions with single hydrophobic substitutions are transported by SMR_Gdx_, but that doubly-substituted guanidiniums are not(Kermani et al., 2020). The SMR_Qacs_ examined, EmrE and integron-associated QacE, did not exhibit transport currents for this series of compounds.

### Kinetics and proton coupling for Gdm^+^ and guanylurea transport

To compare the kinetic properties for transport of guanylurea and the physiological substrate Gdm^+^, we measured the peak amplitudes of the capacitive currents for the four SMR_Gdx_ homologues as a function of substrate concentration. The current amplitudes reflect the initial rate of transport(Bazzone et al., 2017), and their concentration dependence follows Michaelis-Menten kinetics (**Figure 3**, **Table 1**). For all four homologues, the K_m_ value for guanylurea was ∼2-fold lower than that of Gdm^+^. However, the absolute K_m_ values varied over a factor of ∼50 among these proteins. The genomic Gdx-Clo exhibited the lowest K_m_ values (500 μM for Gdm^+^ and 220 μM for guanylurea), and the plasmid-associated Gdx-pAmi exhibited the highest K_m_ values (16 mM for Gdm^+^ and 5 mM for guanylurea). We verified that protein-free liposomes did not exhibit currents at all substrate concentrations tested (**Supplementary** Figure 3).

**Figure 3.**
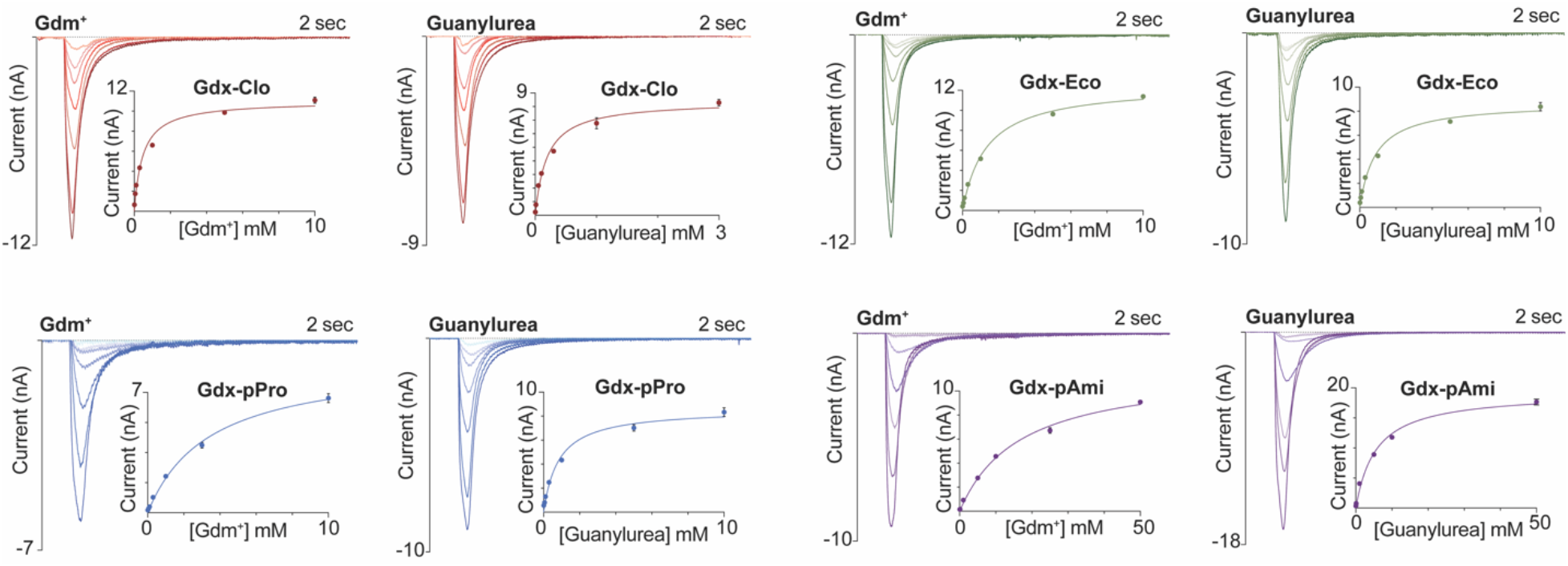
Maximal current amplitudes as a function of Gdm^+^ or guanylurea concentration. Representative transport currents for concentration series of the indicated substrates. Inset: maximum current amplitude as a function of substrate concentration. Solid line represents a fit to the Michaelis-Menten equation. Each of these representative plots was obtained on a single sensor, and error bars represent the SEM for triplicate measurements on that single sensor. K_m_ values reported in Table 1 represent averages from at least 3 independent sensors prepared from 2-3 independent protein preps.

**Table 1.** K_m_ values determined using SSM electrophysiology (pH 7.5)

|  | **Gdm<sup>+</sup> (mM)** | **guanylsurea (mM)** |
| --- | --- | --- |
| Gdx-Clo | $0.5 \pm 0.1$ | $0.22 \pm 0.06$ |
| Gdx-Eco | $1.7 \pm 0.5$ | $0.85 \pm 0.1$ |
| Gdx-pPro | $2.9 \pm 0.5$ | $0.9 \pm 0.2$ |
| Gdx-pAmi | $15.7 \pm 3.3$ | $5.2 \pm 2.0$ |

Our experiments thus far do not allow comparison of transport rates among SMR homologues. As with other measurements that rely on liposome fusion to a lipid bilayer(Stockbridge and Tsai, 2015), the absorption of proteoliposomes to the sensor is subject to considerable variability from experiment to experiment, so that current measurements from different sensors cannot be quantitatively compared(Barthmes et al., 2016). Fusion efficiency can vary from day-to-day, by experimenter, or by sensor batch. In order to normalize maximal currents obtained on different sensors, and thus evaluate differences in transport rate among different proteins (or different mutants of the same protein), we co-reconstituted each SMR_Gdx_ homologue with an internal reference, the fluoride channel Fluc-Bpe(Stockbridge et al., 2015; Stockbridge et al., 2013), so that both the test protein and the reference protein would be absorbed to the sensor in a prescribed molar ratio **(Figure 4A)**. We selected Fluc-Bpe as an internal reference because its extremely high selectivity for fluoride(McIlwain et al., 2021a) prevents cross-reactivity with other substrates or common buffer components. Moreover, its fast fluoride permeation rate and channel mechanism(McIlwain et al., 2021b) yield high sensitivity with small amounts of protein and low concentrations of fluoride. Control experiments with individually reconstituted Fluc-Bpe and SMR_Gdx_ confirm that the SMR_Gdx_ substrates guanidinium and guanylurea do not elicit a response from Fluc-Bpe, and that the SMR_Gdx_ are similarly insensitive to fluoride perfusion (**Figure 4B, Supplementary** Figure 4). By normalizing with respect to the peak fluoride current amplitudes, we obtain good sensor-to-sensor reproducibility (**Figure 4C**). At high protein concentrations or ion fluxes, the maximal currents can be limited by a number of factors such as internal volume, membrane potential, or membrane crowding. However, at the low protein concentrations used in these experiments (2.5 μg Gdx-Clo and 1 μg Fluc-Bpe per mg lipid), the normalized current amplitudes are reasonably linear with respect to the SMR_Gdx_ concentration (**Figure 4D**), indicating that in the concentration regime of these experiments, using Fluc-Bpe as a reference provides a linear readout of transport velocity (**Figure 4D**).

**Figure 4.**
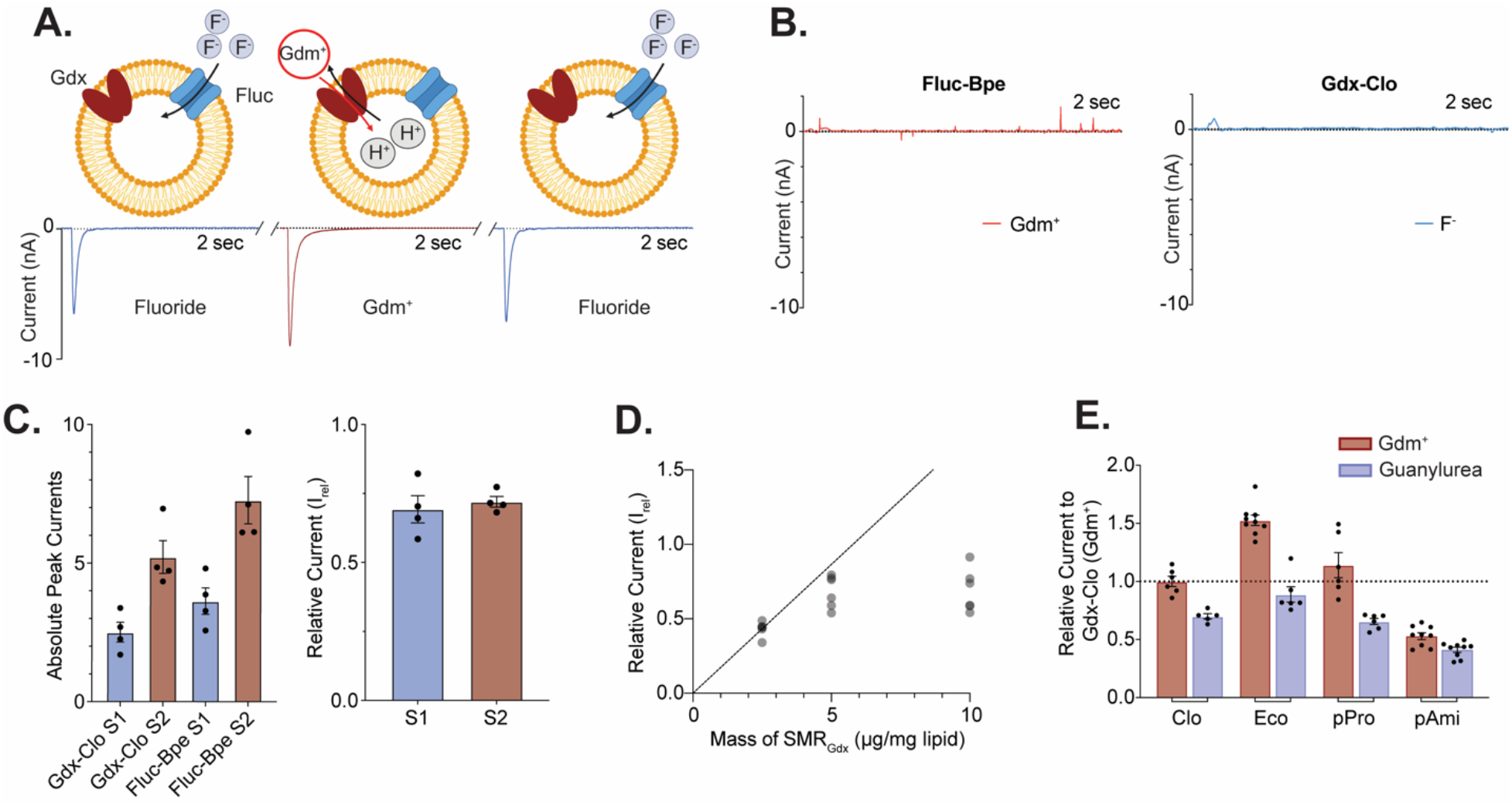
Comparison of maximal velocities for different SMR homologues using an internal reference. A. Schematic showing experimental strategy of co-reconstitution of target transporter and fluoride channel Fluc-Bpe with alternating substrate perfusions. Cartoon made using Biorender. B. Current traces for Gdx-Clo and Fluc-Bpe reconstituted individually do not show substrate cross-reactivity. Tests for cross-reactivity by guanylurea and other SMR_Gdx_ homologues are shown in Supplementary Figure 4. C. Left, peak current amplitude for Gdm^+^ and fluoride currents for two examples of independent sensor preparations. Right, relative Gdm^+^/fluoride current amplitude (I_rel_) for the sensors shown in the left panel. Error bars represent the SEM of individual replicates shown as points. D. I_rel_ as a function of increasing SMR_Gdx_ (Bpe-Fluc held constant at 1 μg/mg lipid). The dashed line represents expected peak current amplitude for a linear response. E. Currents for Gdm^+^ and guanylurea transport by four SMR_Gdx_ homologues normalized against internal Fluc-Bpe reference currents. Each substrate was perfused at a concentration five-fold higher than the K_d_ values measured in Figure 3 in order to compare maximal turnover velocities among the different transporters. Error bars represent the SEM of individual replicates shown as points.

To assess the relative maximal transport velocities of the four SMR_Gdx_ homologues, we evaluated the maximal (initial rate) capacitive currents upon perfusion with substrate at a concentration 5-fold higher than the K_m_ values reported in Figure 3. Using peak fluoride current amplitudes as an internal reference, these experiments show that the transport rates are comparable (within a factor of 3) among the four SMR_Gdx_. For Gdx-Clo, Gdx-Eco, and Gdx-pPro, the maximal velocity for Gdm^+^ is ∼2-fold higher than for guanylurea, whereas for Gdx-pAmi, the turnover rates of guanylurea and Gdm^+^ are comparable (**Figure 4E**).

The negative capacitive currents observed in the SSM electrophysiology experiments presented thus far are in accord with electrogenic transport of >1 H^+^ per substrate, but they do not reveal the precise transport stoichiometry. Prior studies have shown Gdx-Eco possesses a well-coupled 2 H^+^: 1 Gdm^+^ stoichiometry(Kermani et al., 2018; Thomas et al., 2021). However, for SMR_Qac_ EmrE, it has been reported that the transport stoichiometry differs among transported substrates(Robinson et al., 2017). We therefore employed a proteoliposome assay to experimentally assess coupling stoichiometry of Gdx-Clo and plasmid-associated Gdx-pAmi. In these experiments, a 10-fold Gdm^+^ or guanylurea concentration gradient is applied, and the direction of substrate movement is monitored as a function of membrane potential (Fitzgerald et al., 2017; Kermani et al., 2018). When no voltage is applied, substrate is transported down its chemical gradient, coupled to proton efflux. Application of increasingly negative membrane potentials thermodynamically pushes back against the 10-fold substrate gradient; the electrochemical equilibrium point at which no substrate movement occurs is the reversal potential, from which the transport stoichiometry can be calculated using Eq. 2.

In our setup, the membrane potential is established using a potassium gradient and the potassium ionophore valinomycin, and substrate-coupled proton movement is monitored using pyranine, a pH sensitive fluorescent dye, encapsulated inside the liposomes. For coupled 2 H^+^ to 1 substrate transport, electrochemical equilibrium is expected at –60 mV under a 10-fold substrate concentration gradient. We confirmed this value for Gdm^+^ transport by Gdx-Clo (**Figure 5A**), and our experiments show that guanylurea export is carried out with the same stoichiometry by both Gdx-Clo and Gdx-pAmi (**Figure 5B, C**).

**Figure 5.**
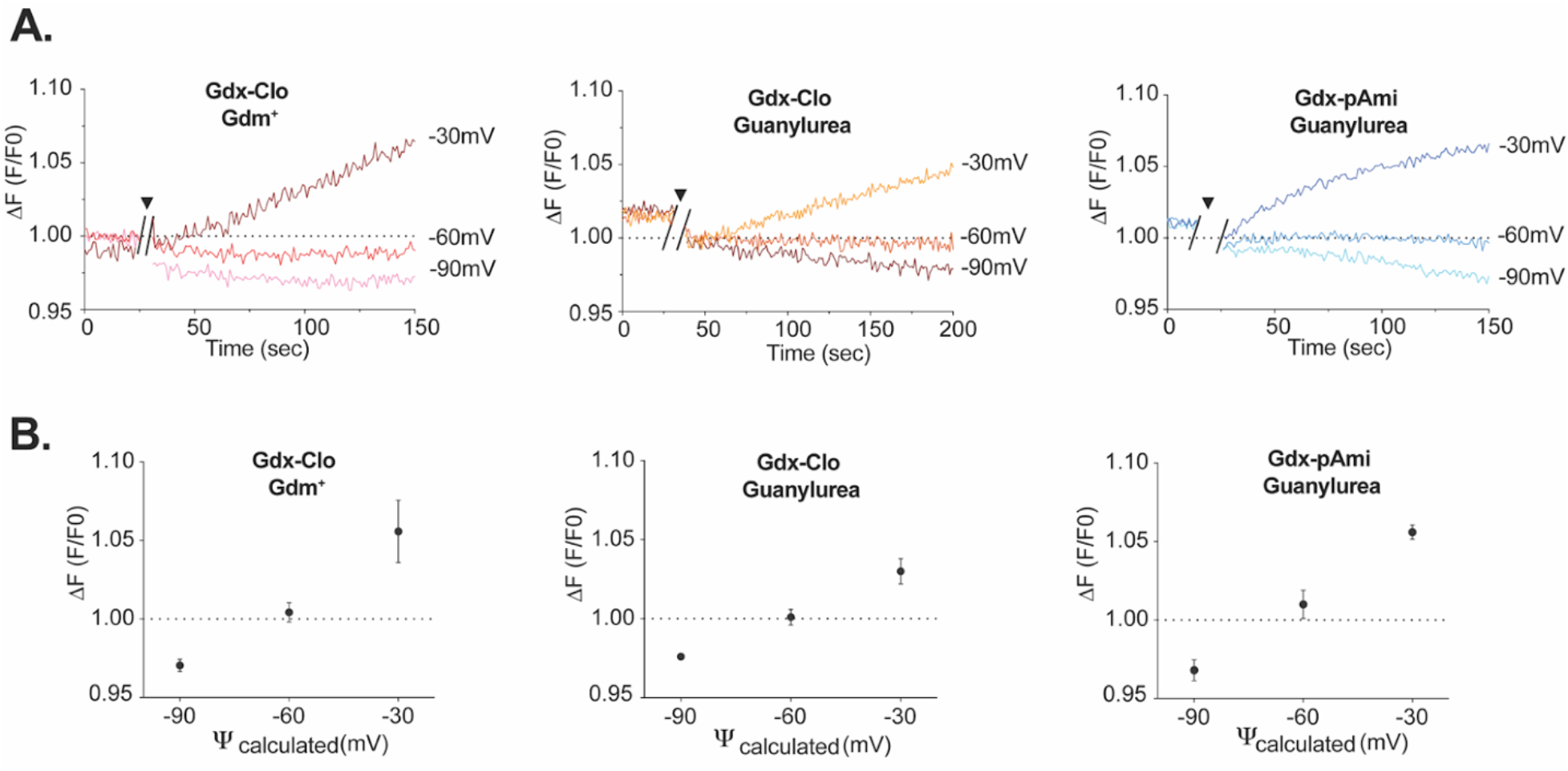
Proton coupling stoichiometry for substrate transport by Gdx-Clo and Gdx-pAmi. A. Change in pyranine fluorescence over time for substrate transport at applied membrane potentials of –30, –60, and –90 mV. After ∼20 seconds of baseline collection, external substrate was added together with valinomycin to establish the 10-fold substrate gradient and membrane potential (indicated by break in trace and triangle). B. Change in pyranine fluorescence as a function of membrane potential for replicate experiments. Error bars represent the SEM for three replicates (–90 and –30 mV) or four replicates (–60 mV). The dashed line represents the equilibrium condition where no proton transport occurs.

### Gdm^+^ and guanylurea binding in Gdx-Clo

To further characterize the pH dependence and thermodynamic properties of Gdm^+^ and guanylurea binding by SMR_Gdx_, we selected the homologue with the best biochemical stability, Gdx-Clo. Although we initially sought to examine substrate binding by Gdx-pAmi as well, the protein requires high salt concentrations for purification and, in detergent, was prone to aggregate over long titrations or at more physiological salt concentrations.

We first exploited intrinsic changes in tryptophan fluorescence to monitor substrate binding at pH values between pH 6 and pH 9 (**Figure 6A**). Gdm^+^ titration induces an increase in tryptophan fluorescence that can be fit with a single site binding isotherm described by Eq. 3 (**Supplementary** Figure 5); separate control experiments showed that binding kinetics are fast and the binding reaction achieved equilibrium prior to measurement. As expected for a model where protons and Gdm^+^ compete for binding to the central glutamates, the apparent binding affinity increases with pH as the central glutamates become increasingly deprotonated (**Table 2**, **Figure 6B**). Although careful NMR experiments with SMR homologue EmrE have shown that the pK_a_ values of the two central glutamates differ(Li et al., 2021; Morrison et al., 2015), the current binding assays do not have the resolution to distinguish independent K_a_ values and report on the averaged behavior of the binding site residues. Using the approximation that the glutamates have equal K_a_ values, the relationship between apparent K_d_ and pH can be fit using Equation 5, yielding an average pK_a_ for the glutamates of 6.7 and a K_d_ for Gdm^+^ of 600 μM.

**Figure 6.**
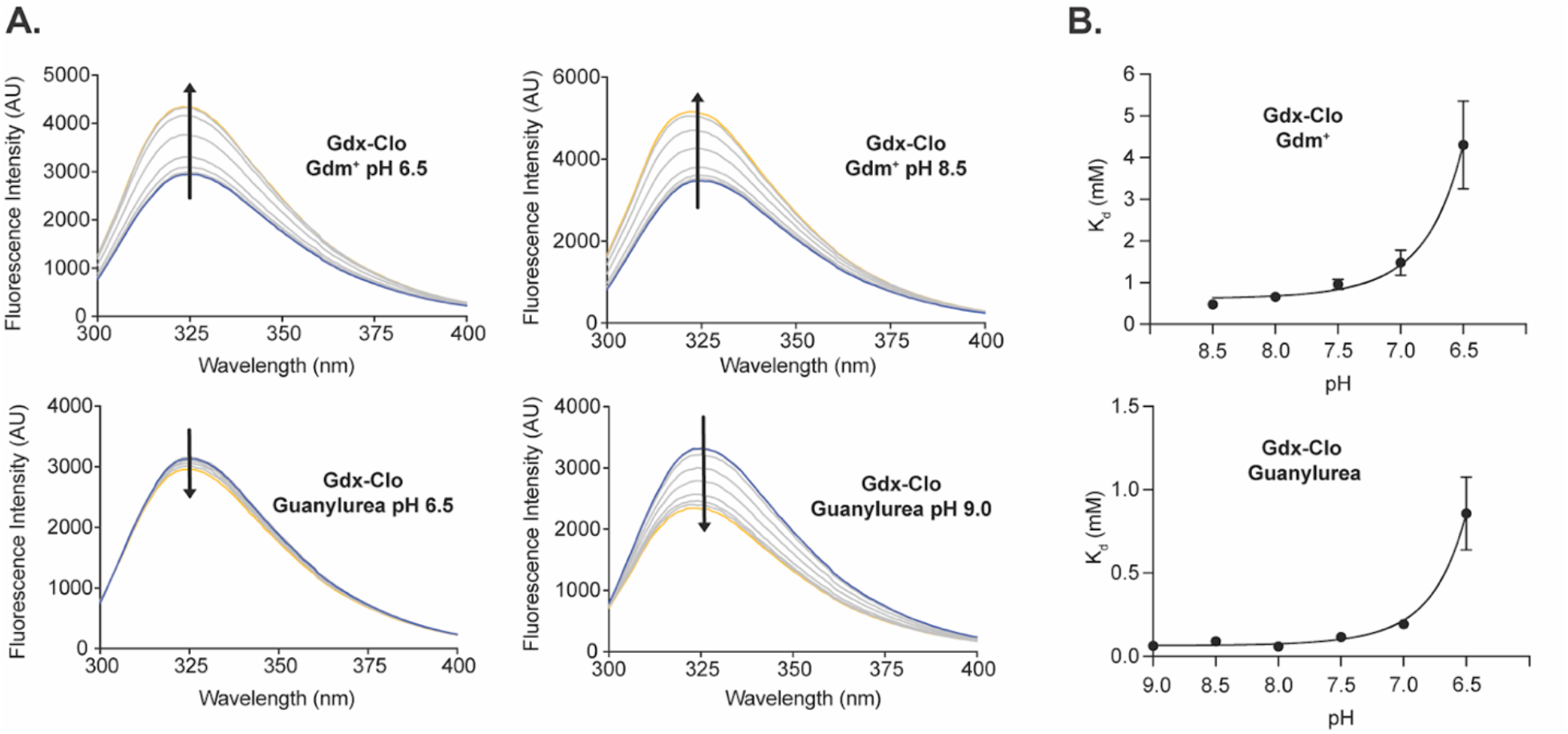
pH dependence of equilibrium substrate binding for Gdx-Clo. A. Tryptophan fluorescence spectra measured at increasing concentrations of Gdm^+^ (top panels) or guanylurea (lower panels) at representative low and high pH values. Arrows denote direction of change in fluorescence intensity with increasing substrate concentration. B. Plot of apparent K_d_ values measured for Gdm^+^ (top) or guanylurea (bottom) as a function of pH. Apparent K_d_ values were determined by fitting tryptophan fluorescence titration isotherms. Fluorescence spectra and fits for all pH values are shown in Supplementary Figure 5. The solid lines represent fits to Equation 5, with a K_d_ value of 600 μM and a pK_a_ of 6.7 for the Gdm^+^ titrations, and a K_d_ value of 70 μM and a pK_a_ of 6.9 for the guanylurea titrations. Error bars represent the SEM of values from 3-4 independent titrations from two independent protein preps.

**Table 2.** K_d_ values for substrate binding to Gdx-Clo as a function of pH.

| **pH** | **Gdm<sup>+</sup> (mM) ±SEM** | **Guanylsurea (mM) ±SEM** |
| --- | --- | --- |
| 6.5 | $4.3 \pm 1.0$ | $0.86 \pm 0.22$ |
| 7.0 | $1.5 \pm 0.3$ | $0.19 \pm 0.02$ |
| 7.5 | $0.96 \pm 0.12$ | $0.12 \pm 0.02$ |
| 8.0 | $0.66 \pm 0.08$ | $0.059 \pm 0.001$ |
| 8.5 | $0.48 \pm 0.10$ | $0.091 \pm 0.016$ |
| 9.0 | Not determined | $0.063 \pm 0.005$ |

Analogous binding experiments were also performed for guanylurea (**Figure 6**, lower panels). In contrast to the tryptophan fluorescence trend observed for Gdm^+^ binding, titration with guanylurea quenched the tryptophan fluorescence signal. Binding data also suggested there was also a low affinity, non-specific component to substrate binding, which became more apparent at high guanylurea concentrations. Fitting the data to a binding model with a linear non-specific component (Eq. 4) yields apparent K_d_ values of the same order as the K_m_ value determined previously. A fit to Eq. 5 indicates that the pK_a_ value of the glutamates is 6.9, in reassuring agreement with the pK_a_ determined in the Gdm^+^ binding experiment, and yields a guanylurea K_d_ of 70 μM. For both substrates, the K_d_ values are similar to the transport K_m_ values, suggesting that the kinetics of substrate binding are fast relative to the conformational change during substrate transport.

Because tryptophan fluorescence is an indirect measurement of binding (made additionally mysterious by the opposite effects of Gdm^+^ and guanylurea on the fluorescence intensity), and to gain additional information on the thermodynamics of substrate binding, we also sought to reproduce our binding measurements using isothermal titration calorimetry (ITC). At pH 8.5, where proton binding to the glutamates is minimized, we observed an exothermic binding reaction for both Gdm^+^ and guanylurea. Gdm^+^ binds with the expected stoichiometry of 1 substrate per protein dimer (**Figure 7A**, **Table 3**). For guanylurea, we also observed a weak endothermic reaction after saturation of the first high-affinity binding site (**Figure 7B**). We did not observe this endothermic contribution in control titrations into buffer, suggesting that it is protein-dependent. We assume that the endothermic reaction represents the non-specific binding observed at high guanylurea concentrations in our tryptophan fluorescence titrations. Subtraction of this component, and setting the binding stoichiometry to one substrate per dimer, yields a fittable binding isotherm. For both substrates, the K_d_ value measured using ITC was in good agreement with the K_d_ value obtained using tryptophan fluorescence, validating the tryptophan fluorescence approach to monitor substrate binding. The ∼4-fold increase in affinity for guanylurea relative to Gdm^+^ was due to increased entropy of the binding reaction. Thermodynamic parameters derived from the ITC data are reported in **Table 3**.

**Figure 7.**
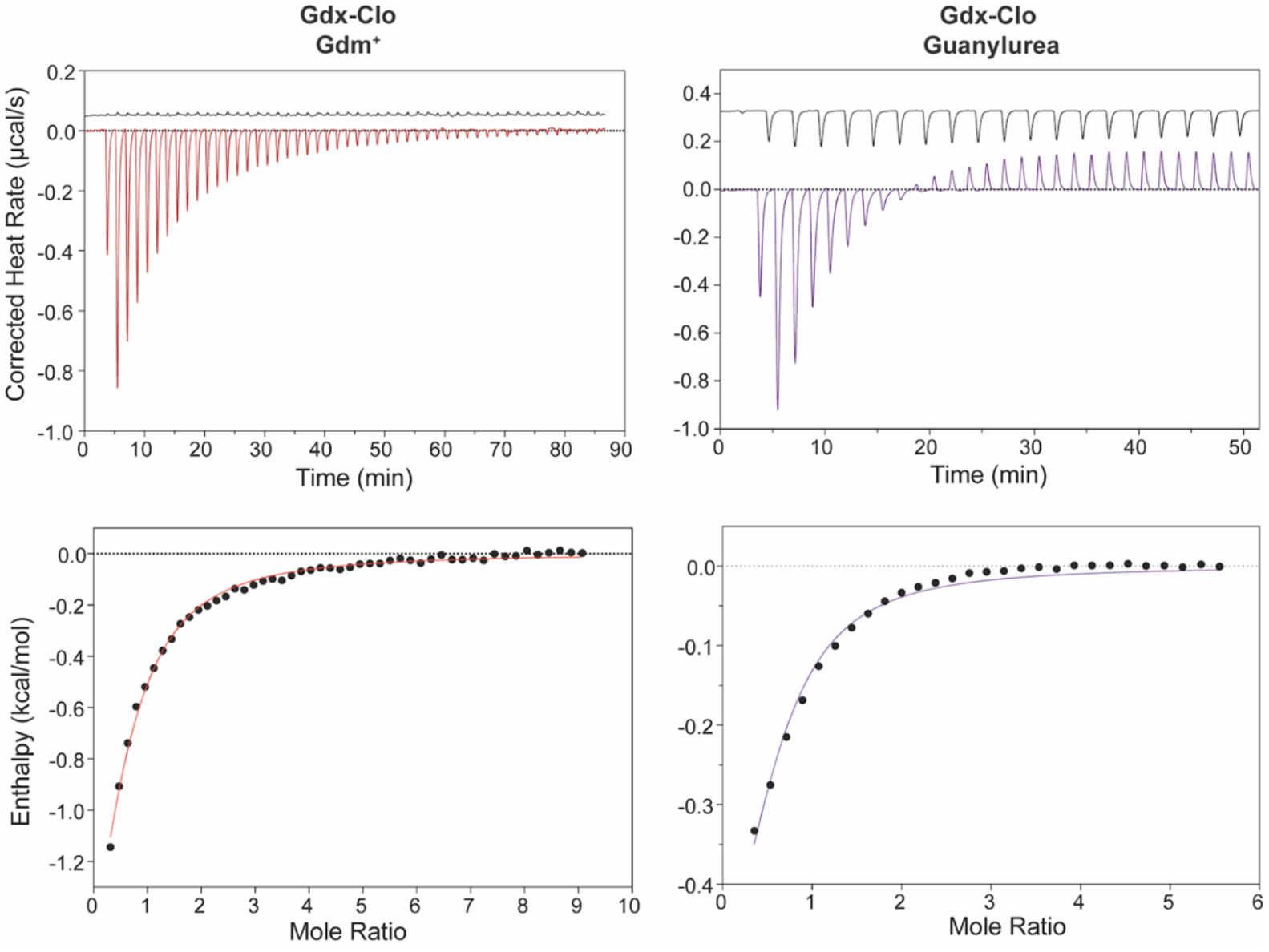
Isothermal titration calorimetry of Gdm^+^ and guanylurea binding to Gdx-Clo. Top panels: Thermograms for Gdm^+^ titrations (left) and guanylurea titrations (right). Lower panels: datapoints show heat absorbed as a function of substrate concentration, fit to equilibrium binding isotherms (solid lines). Equilibrium binding parameters are shown in Table 3.

**Table 3.** Equilibrium binding parameters derived from isothermal titration calorimetry (pH 8.5).

|  | **Gdm<sup>+</sup>** | **Guanylyurea** |
| --- | --- | --- |
| **K<sub>d</sub>** | 490 μM | 230 μM |
| **ΔG** | -4.5 | -5.0 |
| **ΔH** | -3.4 | -3.4 |
| **TΔS** | 1.1 | 1.6 |
| **n** | 0.49 (from fit) | 0.5 (fixed) |

Finally, to determine whether guanylurea occupies the same binding pocket as guanidinium in Gdx-Clo, we solved a crystal structure of Gdx-Clo in the presence of 10 mM guanylurea (**Figure 8**, **Table 4**). Crystals were prepared as in previous studies(Kermani et al., 2022; Kermani et al., 2020), and diffracted to 2.1Å. Two transporters are present in the unit cell, and the maps showed clearly resolved guanylurea density nestled in the binding pocket of one of these transporters (**Figure 8A**). The guanidinium group is poised between the central glutamates, within hydrogen bonding distance, in the same binding mode as observed for phenylguanidinium(Kermani et al., 2020). The carbonyl of guanylurea faces the cleft between helices 2_A_ and 2_B_ (termed the hydrophobic portal(Kermani et al., 2020)), but is just small enough to fit in the binding pocket without requiring a rearrangement of the sidechains lining the portal, in contrast to the slightly larger phenylguanidinium(Kermani et al., 2020). The carbonyl of the guanylurea is twisted slightly out of plane with respect to the guanidinyl group, and is positioned ∼3Å from the electropositive ring edge of portal sidechain F43. There are no other residues within coordination distance of guanylurea, recapitulating the undercoordination of the native substrate Gdm^+^. Other key binding pocket residues (W16, S42, Y59, and W62) contribute to a H-bond network that stabilize the central E13 residues, in the same orientation as seen in other structures (**Figure 8B**)(Kermani et al., 2022; Kermani et al., 2020).

**Figure 8.**
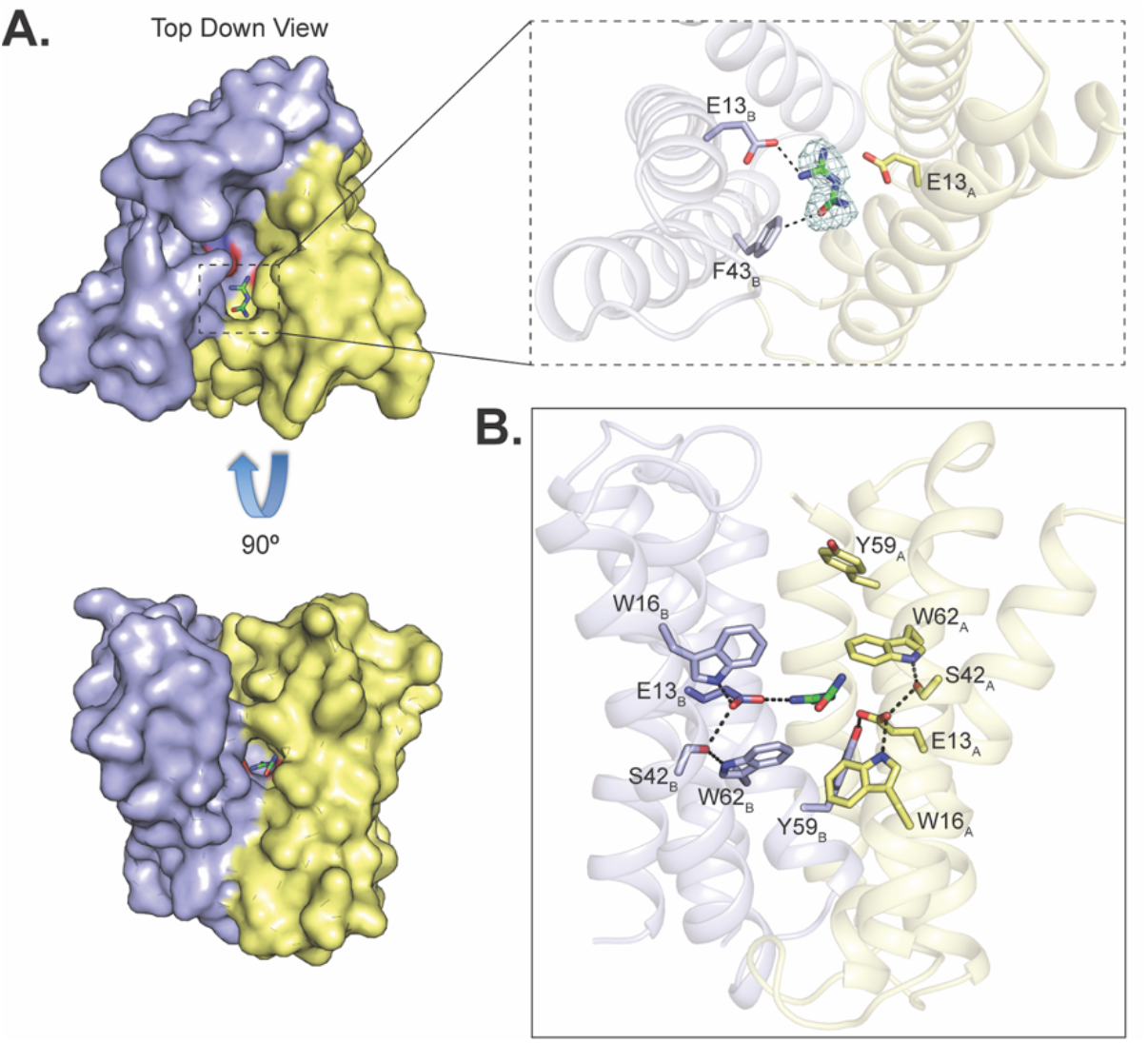
Crystal structure of Gdx-Clo in complex with guanylurea. A. A and B subunits are shown in yellow and blue, respectively, with central glutamates shown as sticks and guanylurea as green sticks. The right upper panel shows the F_o_-F_c_ omit map for guanylurea contoured at 3.5α. B. Putative polar interactions among binding site residues are indicated with dashed lines.

**Table 4.**
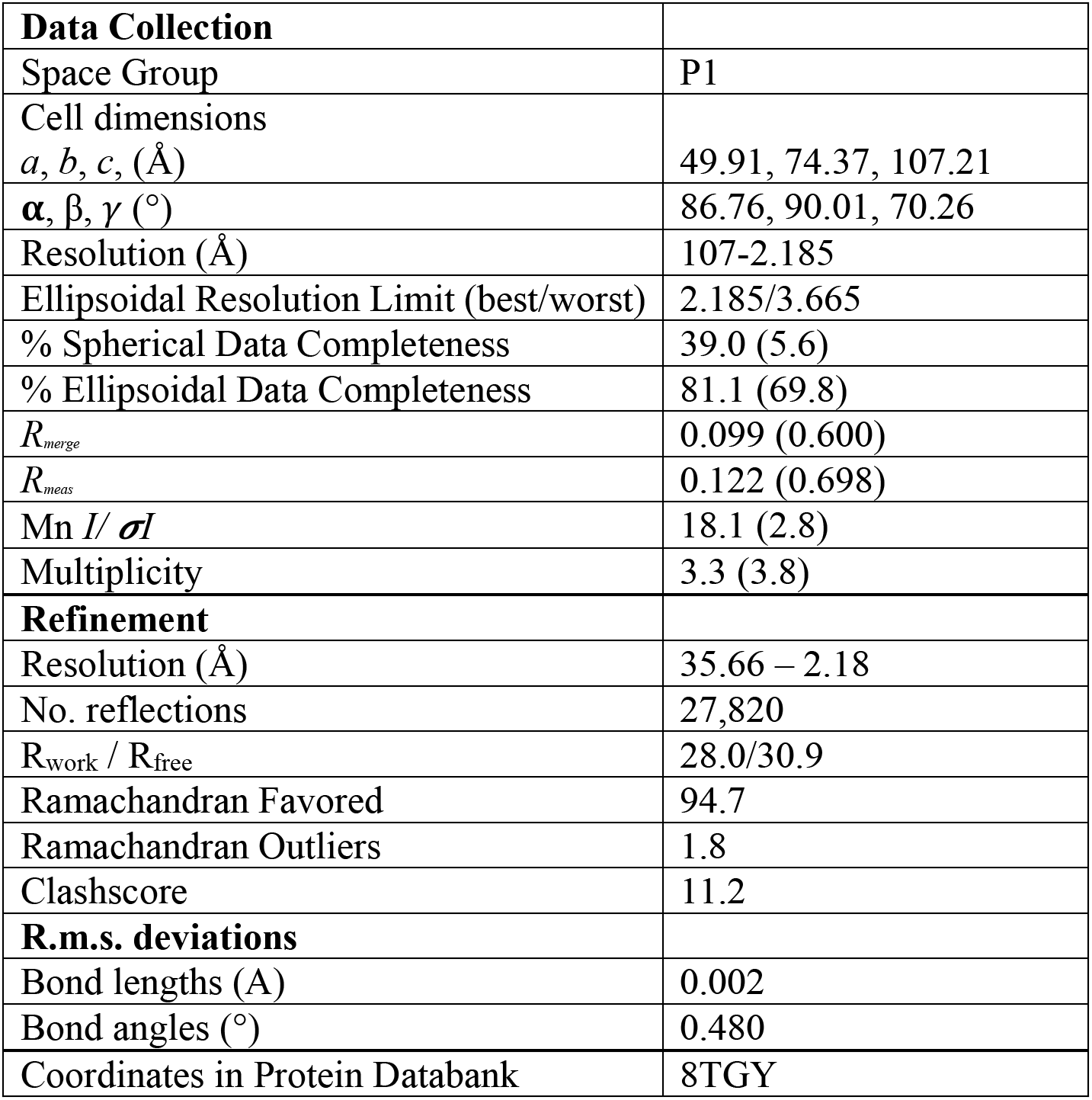
Data collection and refinement statistics for crystal structure of Gdx-Clo in complex with guanylurea.

## Discussion

Microbes are constantly evolving to contend with new environmental pressures, including the recent introduction of anthropogenic chemicals. Major routes for the acquisition of new traits by a microbial population include the gain new genes via HGT transfer events, and the co-option of native proteins’ cryptic functions (functions not under natural selection) to fulfill novel functional roles. Here, we examine a family of transporters, the SMRs, that are associated with both evolutionary processes. In particular, we focus on the SMR_Gdx_, which undergo frequent HGT, despite playing little role in bacterial resistance to classical antimicrobials or antiseptics(Kermani et al., 2020; Slipski et al., 2020). Based on genetic evidence(Li et al., 2023; Martinez-Vaz et al., 2022), we hypothesized a role for the SMR_Gdx_ in transport of metformin metabolites, which structurally resemble the native substrate Gdm^+^, and have accumulated to high levels in waste and surface waters. Our previous work provided preliminary support for this possibility(Martinez-Vaz et al., 2022).

In this study, we investigate whether export of guanylurea or other metformin metabolites is a general property of SMR_Gdx_, and we functionally characterize this activity across multiple plasmid-associated and genomic transporters. We show robust transport of guanylurea, with the same transport stoichiometry, and transport kinetics on the same order as that of the physiological substrate Gdm^+^. Structures of the guanylurea-bound transporter Gdx-Clo show how guanylurea binding exploits the protein’s undercoordination of the native substrate, Gdm^+^(Kermani et al., 2020), fulfilling all of the hydrogen bonds seen for the native substrate without interference from the substrate’s urea group.

It was surprising on its face that the homologue with the most explicit connection to metformin degradation, Gdx-pAmi, had the lowest affinity for guanylurea (5 mM). But for bacteria actively metabolizing metformin as a nitrogen source, very high concentrations of guanylurea are likely to be produced. A prior study measured metformin degradation by an *Aminobacter* culture at a rate of ∼0.7 mM/hour(Li et al., 2023). Considering the culture density and approximating a ∼fL volume for each cell, each bacterium will produce nearly 16 mM internal guanylurea per minute. This back-of-the-envelope calculation illustrates the need for an efflux pathway, and also suggests that bacteria that occupy this niche might be adapted to handle high steady state guanylurea concentrations. It is a truism that an enzyme only needs to be good enough, and apparently, high substrate affinity is not essential for Gdx-pAmi to contribute a selective advantage in the context of metformin degradation.

In summary, this work has functionally characterized an emerging physiological role of the SMR_Gdx_ transporters, export of metformin metabolites. Such a function rationalizes their genetic occurrence with wastewater-associated plasmids and may also have implications for species distribution or horizontal gene transfer in the gut microbiome of patients treated with metformin. Moreover, understanding how bacteria co-opt native physiologies to contend with novel xenobiotics yields insights into microbial adaptation to an increasingly human-impacted biosphere. Our current study highlights a role for active transport in the full microbial degradation pathway for a chemical pollutant, and may inform effective multispecies bioremediation strategies for metformin and other pharmaceuticals in the environment.

## Supplemental Material

**Supplementary Figure 1.**
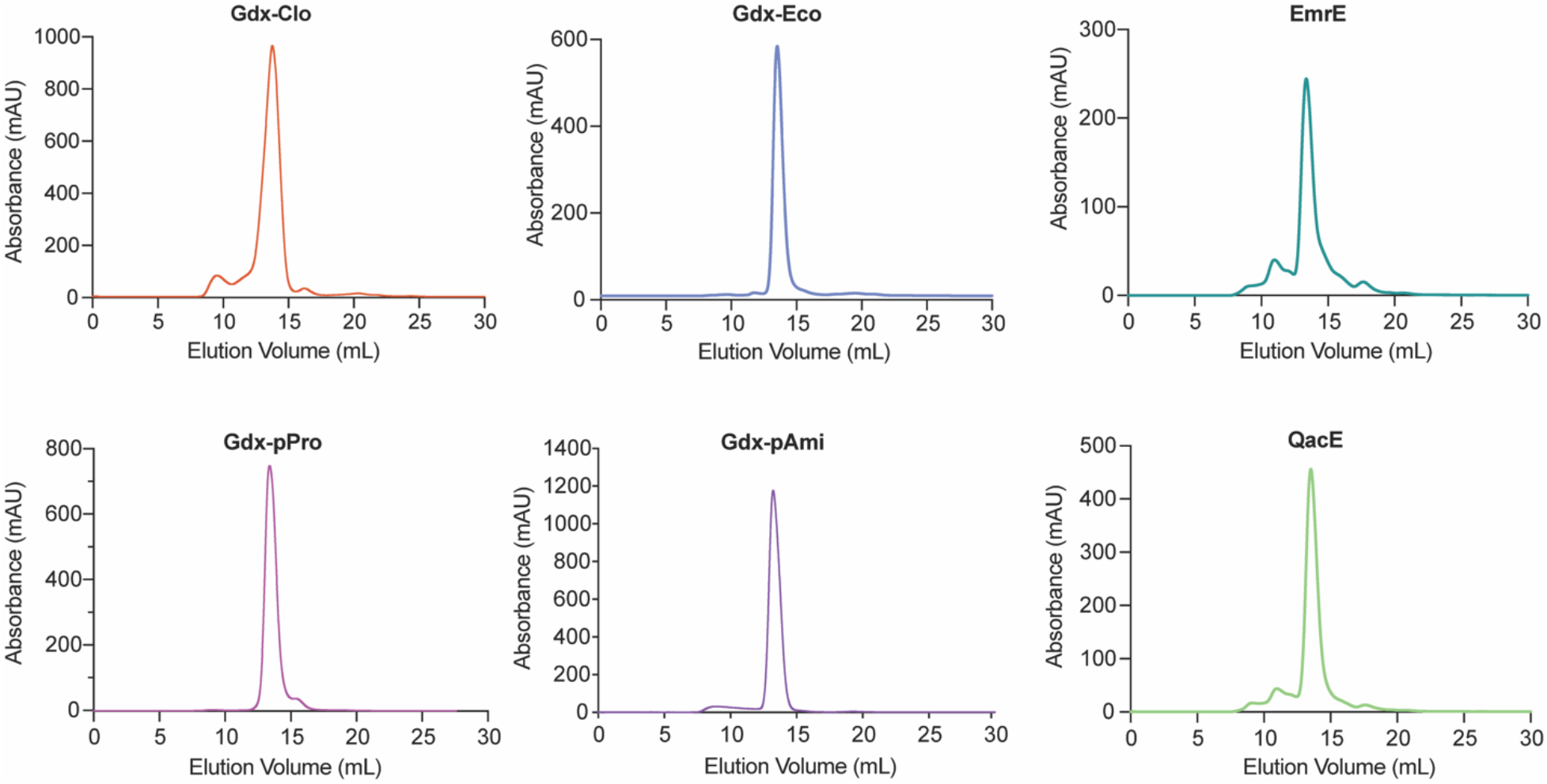
Size exclusion chromatograms for six proteins in this study.

**Supplementary Figure 2.**
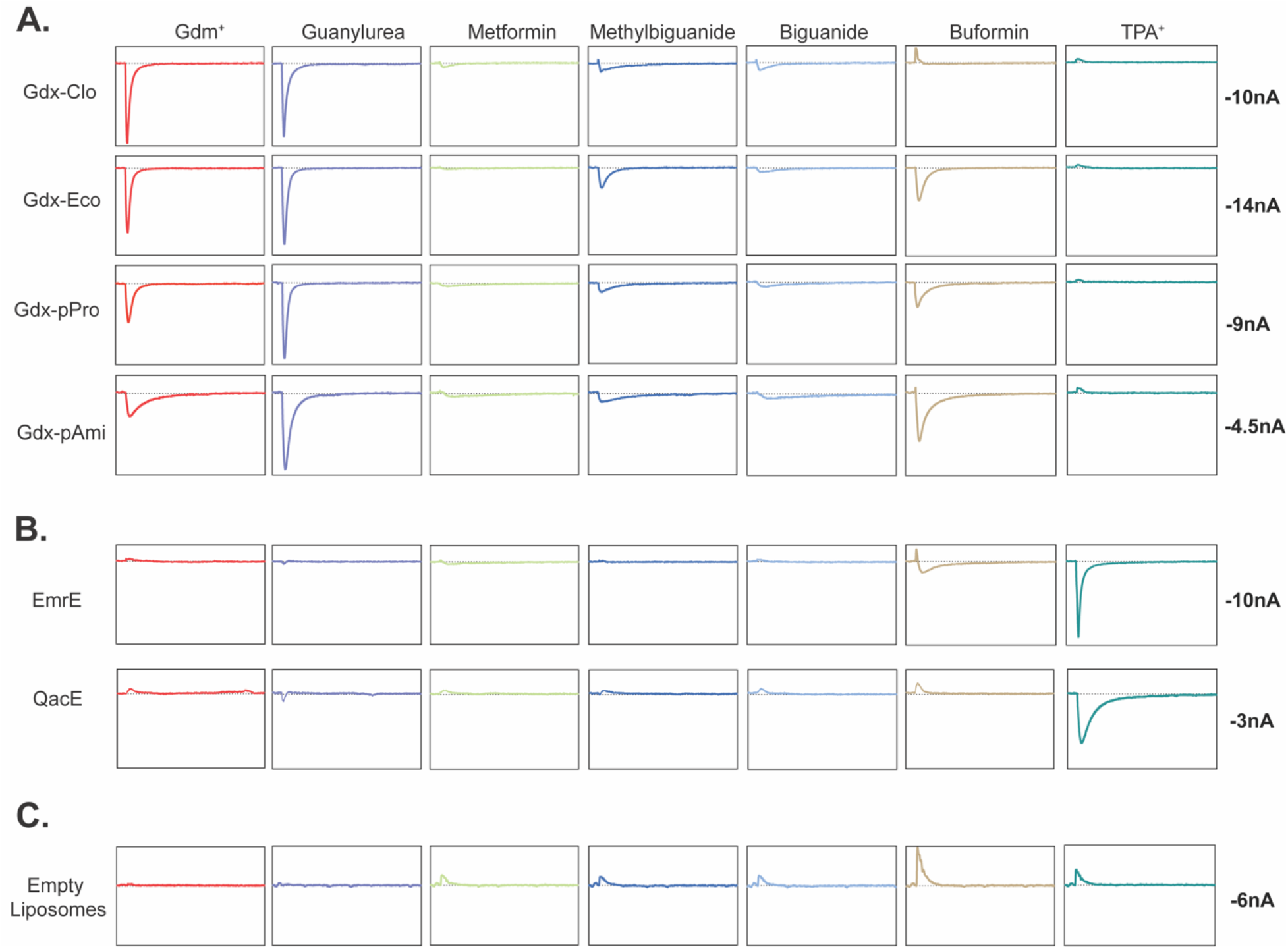
Representative current traces for substrates and transporters summarized in Figure 1, and no-protein controls. Traces for a substrate series are from the same sensor. Box height is equal to current values shown at right.

**Supplementary Figure 3.**
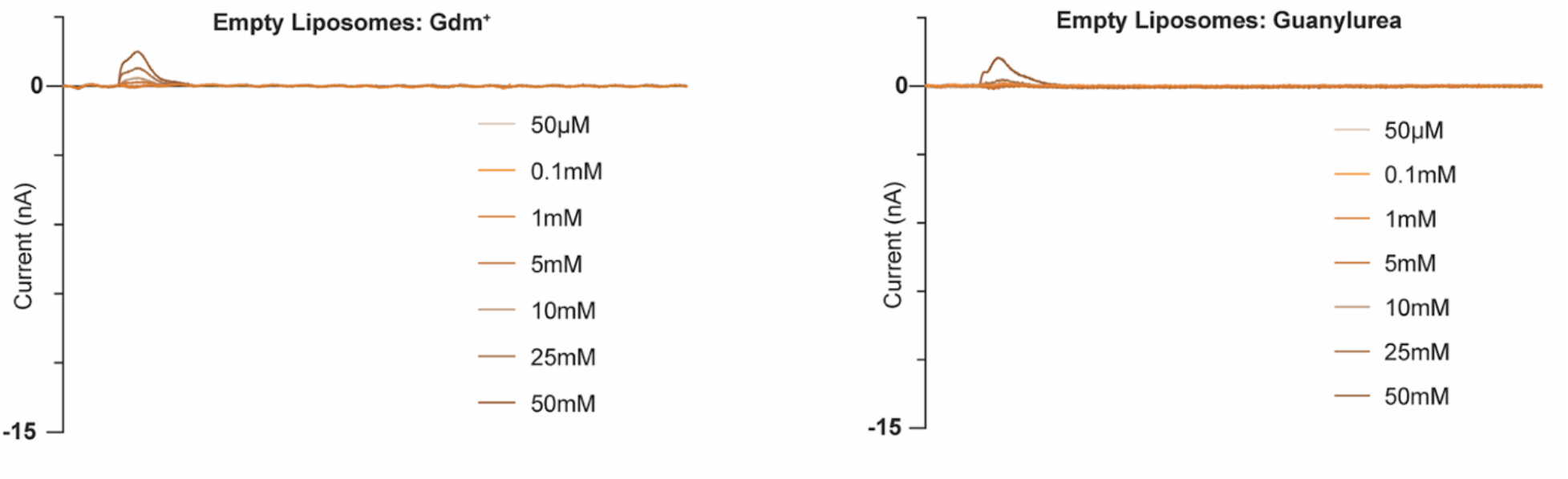
Representative current traces for Gdm^+^ and guanylurea titrations of protein-free liposomes.

**Supplementary Figure 4.**
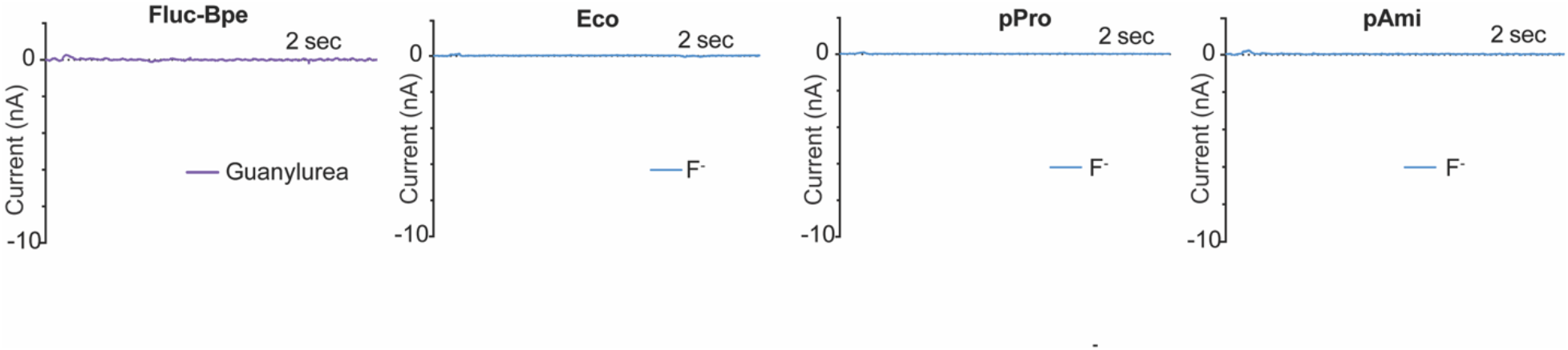
Representative current traces for guanylurea perfusion of Fluc-Bpe and fluoride perfusions of Gdx-Eco, Gdx-pPro, and Gdx-pAmi.

**Supplementary Figure 5.**
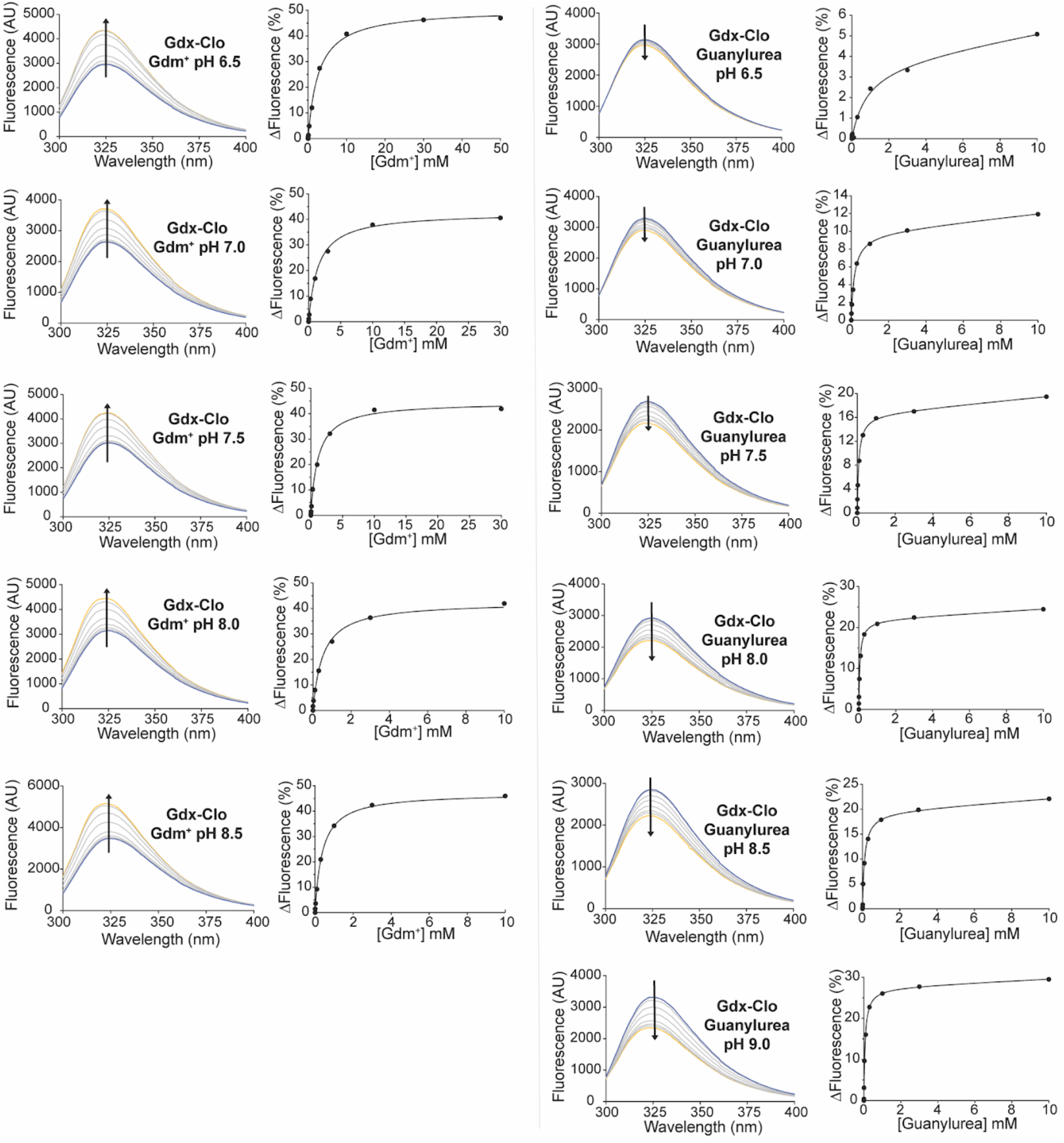
Tryptophan fluorescence spectra and fits to binding isotherms for all data reported in Figure 6 and Table 2.

**Supplementary Table 1.**
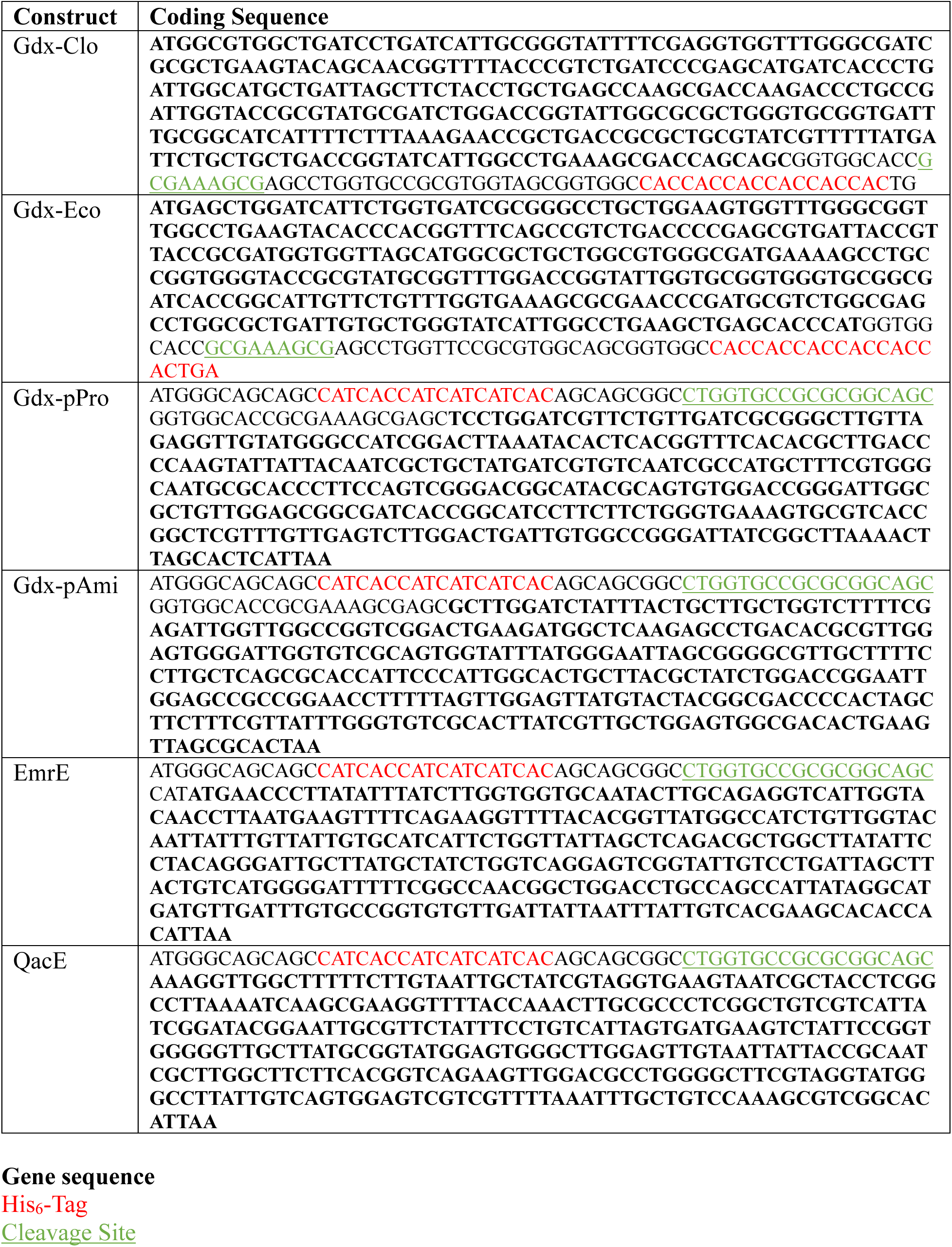
Coding sequences for transporters examined in this study.

## Data Availability

Atomic coordinates for Gdx-Clo bound to guanylurea have been deposited in the Protein Data Bank under accession numbers 8TGY. Source data for figures is available upon request.

## Abbreviations

SMR: Small Multidrug Resistance
Gdx: Guanidinium exporter
Qac: quaternary ammonium cation
Gdm^+^: guanidinium
HGT: horizontal gene transfer
SSM: solid supported membrane
ITC: isothermal titration calorimetry

## Acknowledgements

This work was supported by National Institutes of Health grants R35-GM128768 to RBS and resources of the Advanced Photon Source, a U.S. Department of Energy (DOE) Office of Science User Facility operated for the DOE Office of Science by Argonne National Laboratory under Contract No. DE-AC02-06CH11357. Use of the LS-CAT Sector 21 was supported by the Michigan Economic Development Corporation and the Michigan Technology Tri-Corridor (Grant 085P1000817). The authors declare no competing financial interests.

## Author Contributions

Rachael M. Lucero: Conceptualization, investigation, writing – original draft, visualization; Kemal Demirer: Conceptualization, investigation, validation, writing – original draft, visualization; Trevor Justin Yeh: Conceptualization, investigation, methodology, writing – review and editing, visualization; Randy B. Stockbridge: Conceptualization, investigation, writing – original draft, writing – review and editing, funding acquisition, project administration.

